# Geographic patterns of human allele frequency variation: a variant-centric perspective

**DOI:** 10.1101/2020.07.01.182311

**Authors:** Arjun Biddanda, Daniel P. Rice, John Novembre

## Abstract

A key challenge in human genetics is to describe and understand the distribution of human genetic variation. Often genetic variation is described by showing relationships among populations or individuals, in each case drawing inferences over a large number of variants. Here, we present an alternative representation of human genetic variation that reveals the relative abundance of different allele frequency patterns across populations. This approach allows viewers to easily see several features of human genetic structure: (1) most variants are rare and geographically localized, (2) variants that are common in a single geographic region are more likely to be shared across the globe than to be private to that region, and (3) where two individuals differ, it is most often due to variants that are common globally, regardless of whether the individuals are from the same region or different regions. To guide interpretation of the results, we also apply the visualization to contrasting theoretical scenarios with varying levels of divergence and gene flow. Our variant-centric visualization clarifies the major geographic patterns of human variation and can be used to help correct potential misconceptions about the extent and nature of genetic differentiation among populations.

## Introduction

Understanding human genetic variation, including its origins and its consequences, is one of the long-standing challenges of human biology. A first step is to learn the fundamental aspects of how human genomes vary within and between populations. For example, how often do variants have an allele at high frequency in one narrow region of the world that is absent everywhere else? For answering many applied questions, we need to know how many variants show any particular geographic pattern in their allele frequencies.

In order to answer such questions, one needs to measure the frequencies of many alleles around the world without the ascertainment biases that affect genotyping arrays and other probe-based technologies (International HapMap Consortium 2005; Li et al. 2008). Recent whole-genome sequencing studies (Mallick et al. 2016; Bergström et al. 2019; Fairley et al. 2020) provide these data, and thus present an opportunity for new perspectives on human variation.

However, large genetic data sets present a visualization challenge: how does one show the allele frequency patterns of millions of variants? Plotting a joint site frequency spectrum (SFS) is one approach that efficiently summarizes allele frequencies and can be carried out for data from two or three populations (Gutenkunst et al. 2009). For more than three populations, one must resort to showing multiple combinations of two or three-population SFSs. This representation becomes unwieldy to interpret for more than three populations and cannot represent information about the joint distribution of allele frequencies across all populations. Thus, we need visualizations that intuitively summarize allele frequency variation across several populations.

New visualization techniques also have the potential to improve population genetics education and research. Many commonly used analysis methods, such as principal components analysis (PCA) or admixture analysis, do a poor job of conveying absolute levels of differentiation (McVean 2009; Lawson, van Dorp, and Falush 2018). Observing the genetic clustering of individuals into groups can give a misleading impression of “deep” differentiation between populations, even when the signal comes from subtle allele frequency deviations at a large number of loci (Patterson, Price, and Reich 2006; McVean 2009; Novembre and Peter 2016). Related misconceptions can arise from observing how direct-to-consumer genetic ancestry tests apportion ancestry to broad continental regions. One may mistakenly surmise from the output of these methods that most human alleles must be sharply divided among regional groups, such that each allele is common in one continental region and absent in all others. Similarly, one might mistakenly conclude that two humans from different regions of the world differ mainly due to alleles that are restricted to each region. Such misconceptions can impact researchers and the broader public alike. All of these misconceptions potentially can be avoided with visualizations of population genetic data that make typical allele frequency patterns more transparent.

Here, we develop a new representation of population genetic data and apply it to the New York Genome Center deep coverage sequencing data (see URLs) from the 1000 Genomes Project (1KGP) samples (1000 Genomes Project Consortium et al. 2015). In essence, our approach represents a multi-population joint SFS with coarsely binned allele frequencies. It trades precision in frequency for the ability to show several populations on the same plot. Overall, we aimed to create a visualization that is easily understandable and useful for pedagogy. As we will show, the visualizations reveal with relative ease many known important features of human genetic variation and evolutionary history.

This work follows in the spirit of Rosenberg (2011) who used an earlier dataset of microsatellite variation to create an approachable demonstration of the major features in the geographic distribution of human genetic variation (as well as earlier related papers such as Lewontin 1972; Witherspoon et al. 2007). Our results complement several recent analyses of single-nucleotide variants in whole-genome sequencing data from humans (1000 Genomes Project Consortium et al. 2015; Mallick et al. 2016; Bergström et al. 2019). We label the approach taken here a variant-centric view of human genetic variation, in contrast to representations that focus on individuals or populations and their relative levels of similarity.

### A variant-centric view of human genetic diversity

To introduce the approach, we begin with considering 100 randomly chosen single nucleotide variants sampled from Chromosome 22 of the 1KGP high coverage data (Box 1, Fairley et al. 2020). Figure 1 shows the allele frequency of each variant (rows) in each of the 26 populations of the 1KGP (columns, see Supplemental Table 1 for labels). As a convention throughout this paper, we use darker shades of blue to represent higher allele frequency, and we keep track of the globally minor allele, i.e., the rarer (< 50% frequency) allele within the full sample. The figure shows that variants seem to fall into a few major descriptive categories: variants with alleles that are localized to single populations and rare within them, and variants with alleles that are found across all 26 populations and are common among them.

#### Box 1: Dataset Descriptions and Groupings

We use bi-allelic single nucleotide variants from the New York Genome Center high-coverage sequencing of the 1000 Genomes Project (1KGP) Phase 3 samples (1000 Genomes Project Consortium et al. 2015) (see URLs, Accessed July 22nd, 2019, we include only variants with PASS in the VCF variant filter column). Most of the samples are from an ethnic group in an area (e.g., the “Yoruba of Ibadan,” YRI, or the “Han Chinese from Beijing,” CHB), so the sampling necessarily represents a simplification of the diversity present in any locale (e.g., Beijing is home to several ethnic groups beyond the Han Chinese). For each grouping, the 1KGP typically required each individual to have at least 3 out of 4 grandparents who identified themselves as members of the group being sampled.

The 1KGP further defined five geographical ancestry groups: African (AFR), European (EUR), South Asian (SAS), East Asian (EAS), and Admixed American (AMR). Differing from the 1KGP, we include in the “Admixed in the Americas” (AMR) regional grouping the following populations: “Americans of African Ancestry in SW USA”, “African-Caribbeans in Barbados (ACB)”, and the “Utah Residents (CEPH) with Northern and Western European Ancestry”. We chose this grouping because it is a more straightforward representation of current human geography. We note challenges and caveats of these alternate decisions in the Discussion. Supplemental Table 1 provides a full list of the 26 populations and the grouping into five regions. Figure 7 and Figure S7 provide a complementary view to Figure 2 where the analysis is not based on the five groupings, but instead all 26 populations.

In Figure 5, we present results for variants differing between pairs of individuals from the Simons Genome Diversity Project (SGDP). We include only autosomal biallelic SNVs for variants that pass “filter level 1”, which is the filtering procedure for the majority of analyses used by (Mallick et al. 2016). (see URLs).

In Figure 6 we present results for variants found on five commercially available genotyping arrays: The Affymetrix 6.0 (Affy6) genotyping array, the Affymetrix Human Origins array (HumanOrigins), the Illumina HumanOmniExpress (OmniExpress) array, and the Illumina Omni2.5Exome (Omni2.5Exome), and the Illumina MEGA array (MEGA). We only include autosomal biallelic SNVs in our analysis. Variant lists for each array were downloaded from the manufacturer websites (see URLs).

For assessing the impact of polarizing to ancestral or derived alleles, we downloaded human ancestral allele calls for GRCh38 based on an 8 primate EPO alignment from Ensembl (see URLs). We used only ancestral allele calls supported by at least two outgroup species for our downstream analysis.

**Figure 1.**
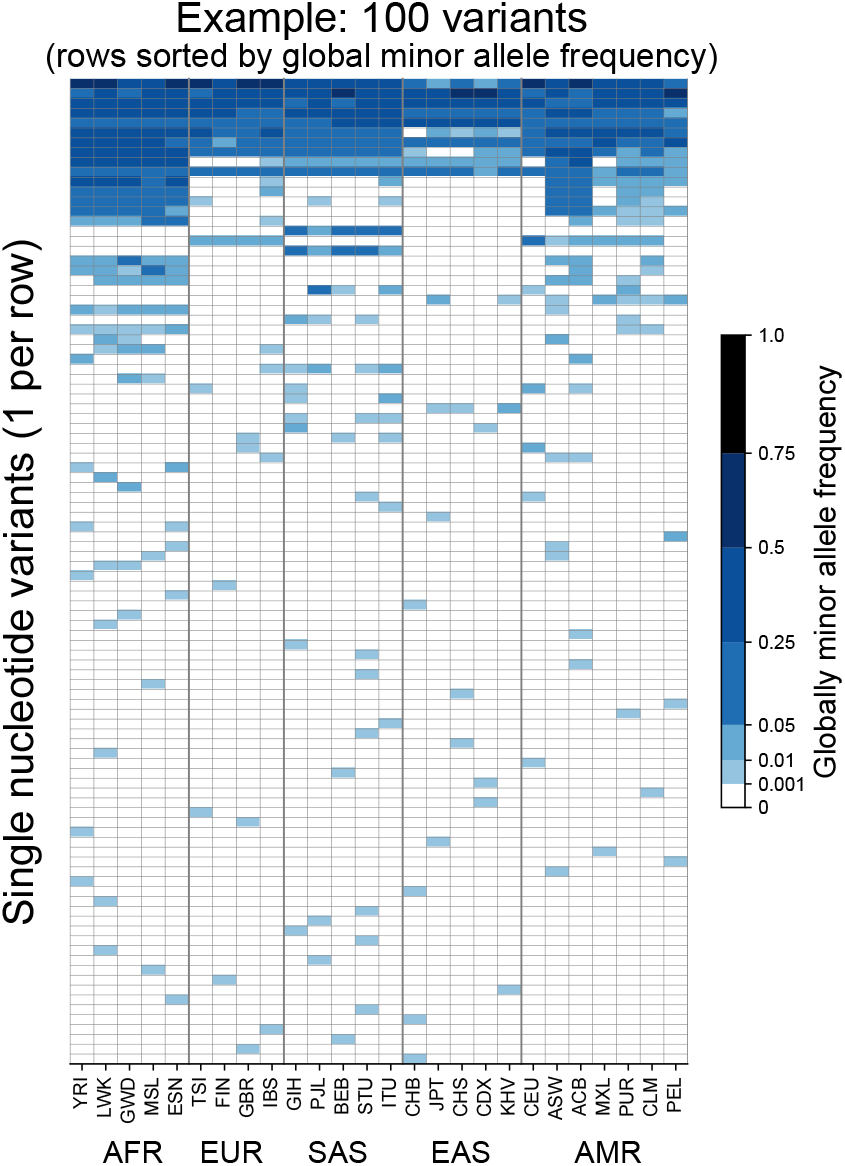
Allele frequencies at 100 randomly chosen variants from Chromosome 22. Frequencies of the globally minor allele across 26 populations from the 1KGP for 100 randomly chosen variants from Chromosome 22. Note that the allele frequency bin spacing is nonlinear to capture variation at low as well as high frequencies.

To investigate whether such patterns hold genome-wide, we devise a scheme that allows us to represent the > 90 million single-nucleotide variants (SNVs) in the genome-wide data (see schematic, Figure 2). First, we follow the 1KGP study in grouping the samples from the 26 populations into five geographical ancestry groups: African (AFR), European (EUR), South Asian (SAS), East Asian (EAS), and Admixed American (AMR) (Figure 2A, Box 1). For clarity, we modify the original 1KGP groupings slightly for this project (by including several samples from the Americas in the AMR grouping, see Box 1). While human population structure can be dissected at much finer scales than these groups (e.g., Leslie et al. 2015; Novembre and Peter 2016), the regional groupings we use are a practical and instructive starting point—as we will show, several key features of human evolutionary history become apparent, and many misconceptions about human differentiation can be addressed efficiently with this coarse approach (see Discussion). As any such groupings are necessarily arbitrary, we also show results without using regional groupings to calculate frequencies (see section Finer-scale resolution of variant distributions below).

**Figure 2.**
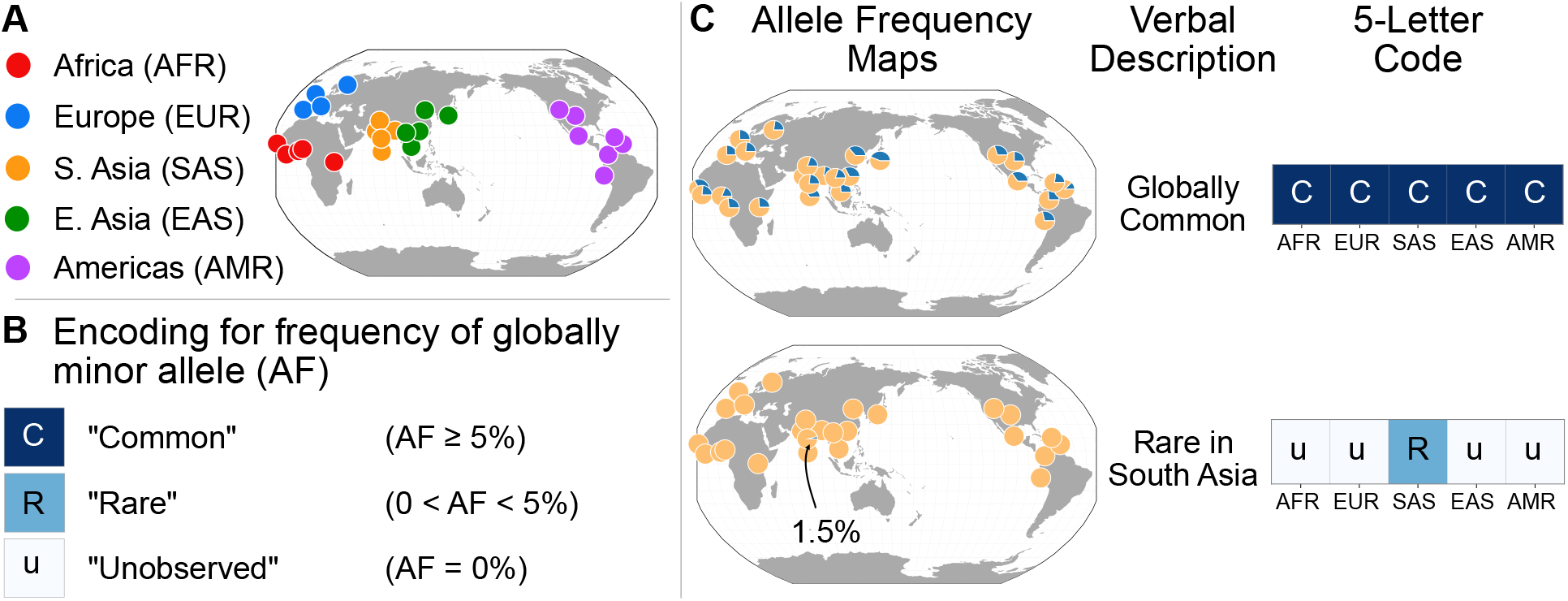
A simple coding system to represent geographic distributions of variants. **A:** Regional groupings of the 26 populations in the 1KGP Project. **B:** Legend for minor allele frequency bins. **C:** Two examples of how a verbal description of an allele frequency map can be communicated equivalently with a 5-letter code (yellow signifies the major allele frequency, blue signifies the minor allele frequency in the pie charts).

To represent the geographic distributions of alleles compactly, we give every variant a five-letter code according to its allele frequencies across regions (Figure 2A). More precisely, for each bi-allelic single nucleotide variant, we identify the global rarer (minor) allele. Then for each region, we code the allele’s frequency as ‘u’, ‘R’, or ‘C’, based on whether the allele is “(undetected,” “(R)are,” or “(C)ommon” (Figure 2B). Finally, we concatenate the allele’s regional frequency codes in the fixed (and arbitrary) order: AFR, EUR, SAS, EAS, and AMR. This procedure generates a “geographic distribution code” for each variant. For example, the code ‘CCCCC’ represents a variant that is common across every region, while ‘uuRuu’ represents a variant that is rare in South Asia and unobserved elsewhere (Figure 2C).

This scheme requires a few choices. To distinguish between “rare” and “common” alleles, we used a threshold of 5% frequency. For comparison, we also show results using a 1% frequency threshold (Figure S1A). For 96.6% of variants in the dataset with high-quality ancestral allele calls (Box 1), the globally minor allele is the derived (younger) allele, and for comparison we also produced results tracking the derived rather than the globally minor allele (Figure S1C). Neither changing the frequency threshold to 1% nor tracking the derived allele meaningfully affects the basic observations that follow.

Next, we coded all ~92 million biallelic SNVs in the dataset and tabulated the proportions of each geographic distribution code. We display the codes in a vertical stack from the most abundant code at the bottom to the least abundant at the top with the height of each code proportional to its abundance, so that the cumulative proportions of the rank-ordered codes are easily readable (Figure 3).

**Figure 3.**
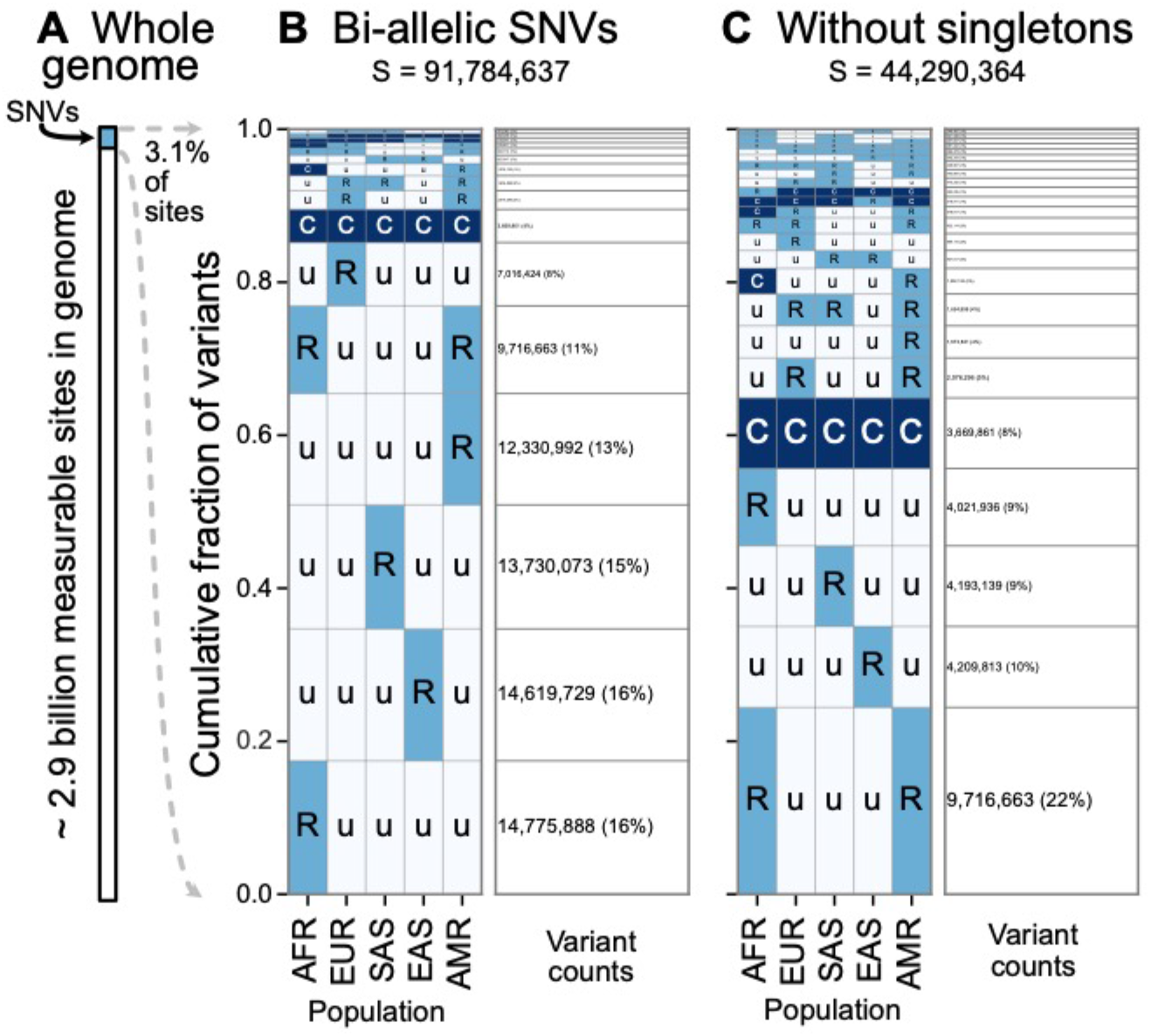
A summary of geographic distributions in human SNVs. **A:** We observe variants at ~3.1% of the measurable sites in the reference human genome (GRCh38). A measurable site is one at which it is possible to detect variation with current sequencing technologies (currently approximately 2.9 Gb out of 3.1 Gb in the human genome; see URLs). **B and C:** The relative abundance of different geographic distributions for 1KGP variants, (B) including singletons, and (C) excluding singletons. In panels B and C, the right-hand rectangles show the number and percentage of variants that fall within the corresponding geographic code on the left-hand side; distribution patterns are sorted by their abundance, from bottom-to-top. See Figure 2 for an explanation of the 5-letter ‘u’,’R’,’C’ codes. The proportion of the genome with variants that have a given geographic distribution code can be calculated from the data above (for example, with the ‘Ruuuu’ code, as 17% × 3.1% = 0.53%).

The distribution of codes is heavily concentrated, with 85% of variants falling into just eight codes out of the 242 (35 – 1) that are possible. Of the top eight codes, the top four codes represent rare variants that are localized in a single region. The fifth most abundant code, ‘RuuuR’, represents rare variants found in Africa and the Admixed Americas (which includes African American individuals, for example). The sixth code is another set of localized rare variants (‘uRuuu’, i.e., variants rare in EUR). The seventh code is ‘CCCCC’ or “globally common variants.” The eighth most abundant category, ‘uRuuR’, represents rare variants found in Europe and the Admixed Americas. Conspicuously infrequent in the distribution are variants that are common in only one region outside of Africa and absent in others (e.g., ‘uCuuu’, ‘uuCuu’, ‘uuuCu’, ‘uuuuC’). Instead, when a variant is found to be common (>5% allele frequency) in one population, the modal pattern (37.3%) is that it is common across the five regions (‘CCCCC’). Further, 63% of variants common in at least one region are also globally widespread, in the sense of being found across all five regions. This number rises to 82% for variants common in at least one region outside of Africa (Figure S2 and S3).

Singleton variants—alleles found in a single individual—are the most abundant type of variant in human genetic data and are necessarily found in just one geographic region. To focus on the distributions of non-singleton variants, we removed singletons and retallied the relative abundance of patterns (Figure 3C). Removing singletons reduces the absolute number of variants observed by 48.2% (91,784,637 vs. 44,290,364). Without singletons, we see more clearly the abundance of patterns that have rare variants shared between two or more regions (codes with two ‘R’s and one ‘u’, such as ‘uuRRu’ or ‘RRuuu’).

The patterns observed here are interpretable in light of some basic principles of population genetics. Rare variants are typically the result of recent mutations (Mathieson and McVean 2014; Kiezun et al. 2013; Kimura and Ohta 1973; Albers and McVean 2020). Thus, we interpret the localized rare variants (such as ‘Ruuuu’ or ‘uuuRu’) as mostly young mutations that have not had time to spread geographically. The code ‘CCCCC’ (globally common variants), likely comprises mostly older variants that arose in Africa and were spread globally during the Out-of-Africa diaspora and other dispersal events (see Box 2). The appearance of rare variants shared between two or more regions (codes with two ‘R’s and three ‘u’s, such as ‘uuRRu’ or ‘RRuuu’) is likely the signature of recent gene flow between those regions (Box 2) (Platt et al. 2019; Mathieson and McVean 2014; Gutenkunst et al. 2009). In particular, the abundant ‘RuuuR’ and ‘uRuuR’ codes likely represent young variants that are shared between the Admixed Americas and Africa (‘RuuuR’) or Europe (‘uRuuR’) because of the population movements during the last 500 years that began with European colonization of the Americas and the subsequent slave trade from Africa. We interpret the 10th most abundant code (‘CuuuR’, Figure 3B) as mostly variants that were lost in the Out-of-Africa bottleneck and subsequently carried to the Americas by African ancestors. There is a relative absence of variants that are common in only one region outside of Africa and absent across all others (e.g., ‘uCuuu’, ‘uuCuu’, ‘uuuCu’, ‘uuuuC’). This is consistent with human populations having not diverged deeply, in the sense that there has not been sufficient time for genetic drift to greatly shift allele frequencies among them (Box 2). To help make this clear, consider the alternative scenario—in a deep, multiregional origins model (Wolpoff, Wu, and Thorne 1984), one would expect many more variants to be common to one region and absent in others (‘uCuuu’ or ‘uuuCu’ for example, see Box 2). Overall, these results reflect a timescale of divergence consistent with the Recent-African-Origin model of human evolution as well as subsequent gene flow among regions (Cann, Stoneking, and Wilson 1987; Stringer and Andrews 1988; Thomson et al. 2000; Ramachandran et al. 2005; Pickrell and Reich 2014).

#### Box 2: Theoretical Modeling

We can use theoretical models to estimate what our visualizations would look like for two populations in simple contrasting cases of “deep” divergence, “shallow” divergence, and “shallow” divergence with gene flow. The shallow case is calibrated to be qualitatively consistent with the Recent-African-Origin model with subsequent gene flow. The deep case mimics a multiregional model of human evolution (Wolpoff, Wu, and Thorne 1984). For each case, we computed the expected abundances of distribution codes in a simple model of population divergence: two modern populations of *N* individuals each that diverged *T* generations ago from a common population of *N* individuals (see Appendix A for information about this calculation). We model gene flow by including recent admixture: individuals in Population A derive an average fraction *α* of their ancestry from Population B and vice versa. This simplified model neglects many of the complications of human population history, including population growth, continuous historical migration, and natural selection, but it captures the key features of common origins, divergence, and subsequent contact.

In this model, the key control parameter is *T/2N*, the population-scaled divergence time. Human pairwise nucleotide diversity (~1 × 10^-3^) and per-base-pair per-generation mutation rate (~1.25 × 10^-8^) imply a Wright-Fisher effective population size of *N* = 2 × 10^4^ individuals. The Out-of-Africa divergence is estimated to have occurred approximately 60,000 years ago (Nielsen et al. 2017). Assuming a 30-year generation time (Fenner 2005) gives *T/2N* = 0.05. We compare this scenario with *T/2N* = 0.5, corresponding to a deeper divergence of approximately 600,000 years ago.

Figure 4A shows the expected patterns in a sample of 100 individuals from each population for deep divergence (*T/2N* = 0.5), shallow divergence (*T/2N* = 0.05) without admixture, and shallow divergence with admixture (*α* = 0.02). The shallow divergence model with or without admixture reproduces the preponderance of ‘Ru’ and ‘CC’ mutations seen in the data, while the deep divergence model shows many more ‘Cu’ and many fewer ‘CC’ mutations. The case with admixture shows a slight increase in variant sharing (‘RR’ alleles increase from 1.3% of variants to 4.2%; ‘RC’ and ‘CR’ alleles increase from 6% to 10%; ‘CC’ alleles comprise 23% in both cases).

**Figure 4.**
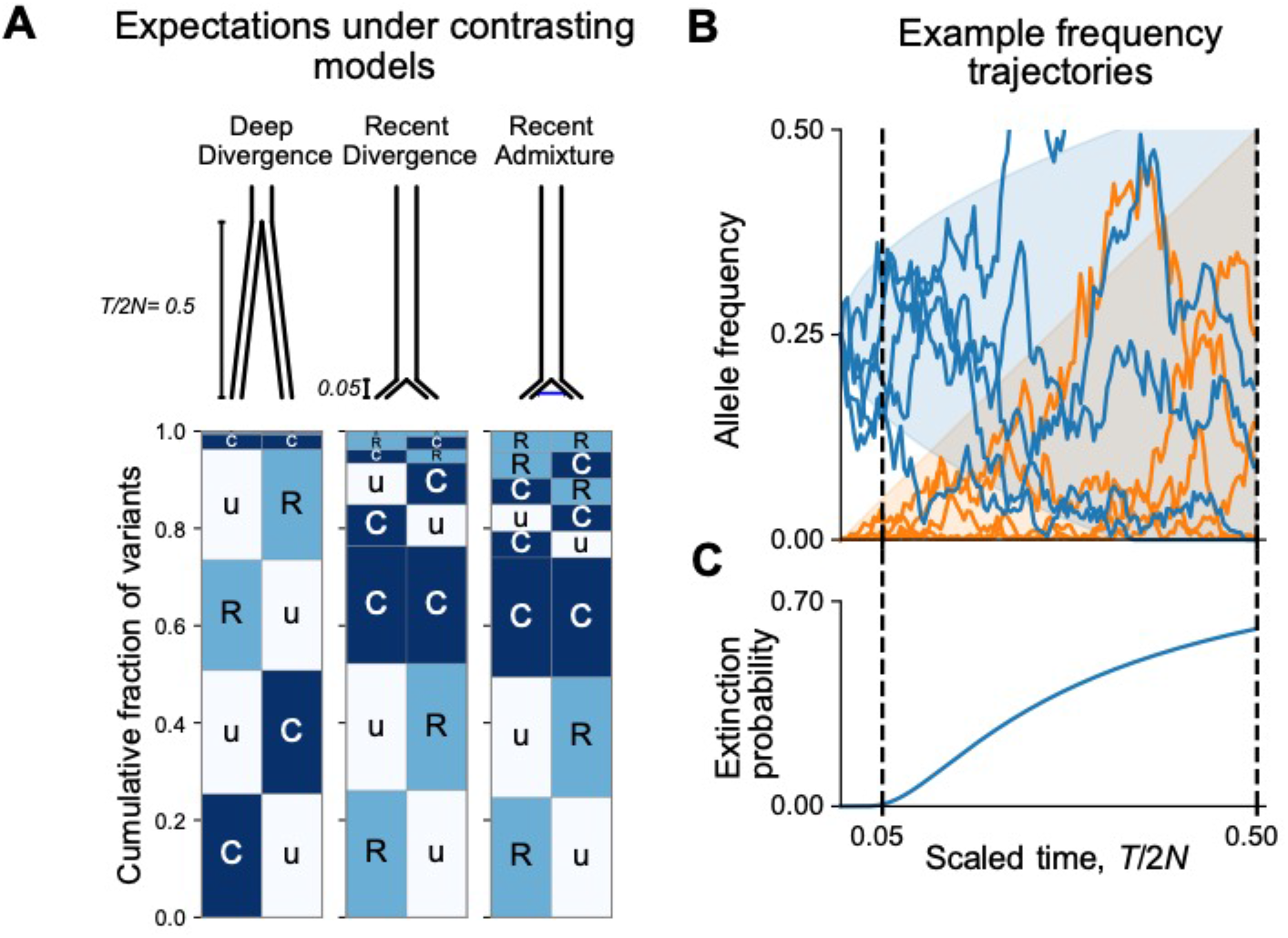
Allele frequency patterns depend on the time since population divergence and levels of admixture. **A:** Expected geographic distribution code abundances in a sample of 100 diploid individuals from each of two populations, for deep divergence (*T/2N* = 0.5, *α* = 0), recent divergence without admixture (*T/2N* = 0.05, *α* = 0), and recent divergence with admixture (*T/2N* = 0.5, *α* = 0.02). **B:** Simulated allele frequency time series for mutations starting at 25% frequency (blue) and new mutations entering the population since the split (orange). **C:** The probability of extinction of a mutation starting at 25% frequency (see Appendix B).

We can understand the relationship between the split time and geographic distribution abundances heuristically as follows. During an interval of *Δt* generations, the frequency of a neutral mutation starting at frequency *f* changes randomly by a typical amount 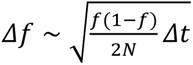. Consider a mutation that is at 25% frequency, i.e., common, in the ancestral population at the time of the split (Figure 4B). At time Δt/2N = 0.05 after the split, the frequency of the mutation is likely to be in the interval (15%, 35%) in both populations and will be assigned the code ‘CC’. On the other hand, by time *Δt/2N* = 0.5 after the split, the mutation has a significant chance of going extinct in one or both populations (Figure 4C). Mutations that go extinct in one population but not the other will typically be assigned a code ‘Cu’ or ‘uC’.

At the same time, new mutations are constantly entering the evolving populations. These new mutations will be private to one population (‘Ru’ or ‘Cu’) and the overwhelming majority will go extinct before reaching detectable frequencies. Conditional on non-extinction, the expected frequency of a neutral mutation increases linearly with time (see Appendix B). As a result, the frequencies of new mutations since the split time *Δt* will mostly be contained in a triangular envelope (*f* < *Δt/2N* (Figure 4B). For recent divergence, the new mutations will be assigned code ‘Ru’ or ‘uR’, while in deeply diverged populations they may be categorized as ‘Cu’ or ‘uC’.

### The variants that differ between a pair of individuals

While Figure 3 illustrates genetic variants found in a large, global sampling of human diversity, it does not show what to expect for the variants that differ between pairs of individuals. Are the variants that differ between two individuals more often geographically widespread or spatially localized?

To address this question, we considered the variants carried by pairs of individuals from the whole-genome sequencing data of the Simons Genome Diversity Project (SGDP) (Mallick et al. 2016) (Figure 5). The SGDP sampled 300 individuals from 142 diverse populations. We use the SGDP data to avoid ascertainment biases that might arise from looking at individuals within the same dataset we use to measure allele frequencies. Figure 5 shows a representative subset with 6 pairs chosen from 3 populations (Figure S6, shows a larger set of examples). For each pair we see some variants that were undiscovered in the 1KGP data (denoted *S_u_* in the figure). These account for 17–20% of each set of pairwise SNVs and are likely rare variants. We see that the variants that differ between each pair of individuals are typically globally widespread (i.e., codes with no ‘u’s, with proportions out of the total S varying from 54%–76% for the pairs in Figure 5.) The observation of mostly globally common variants in pairwise comparisons may seem counterintuitive considering the abundance of rare, localized variants overall. However, precisely because rare variants are rare, they are not often carried by either individual in a pair. Instead, pairs of individuals mostly differ because one of them carries a common variant that the other does not; and as Figure 3 already showed, common variants in any single location are often common throughout the world (also see Figures 7 and S1).

**Figure 5.**
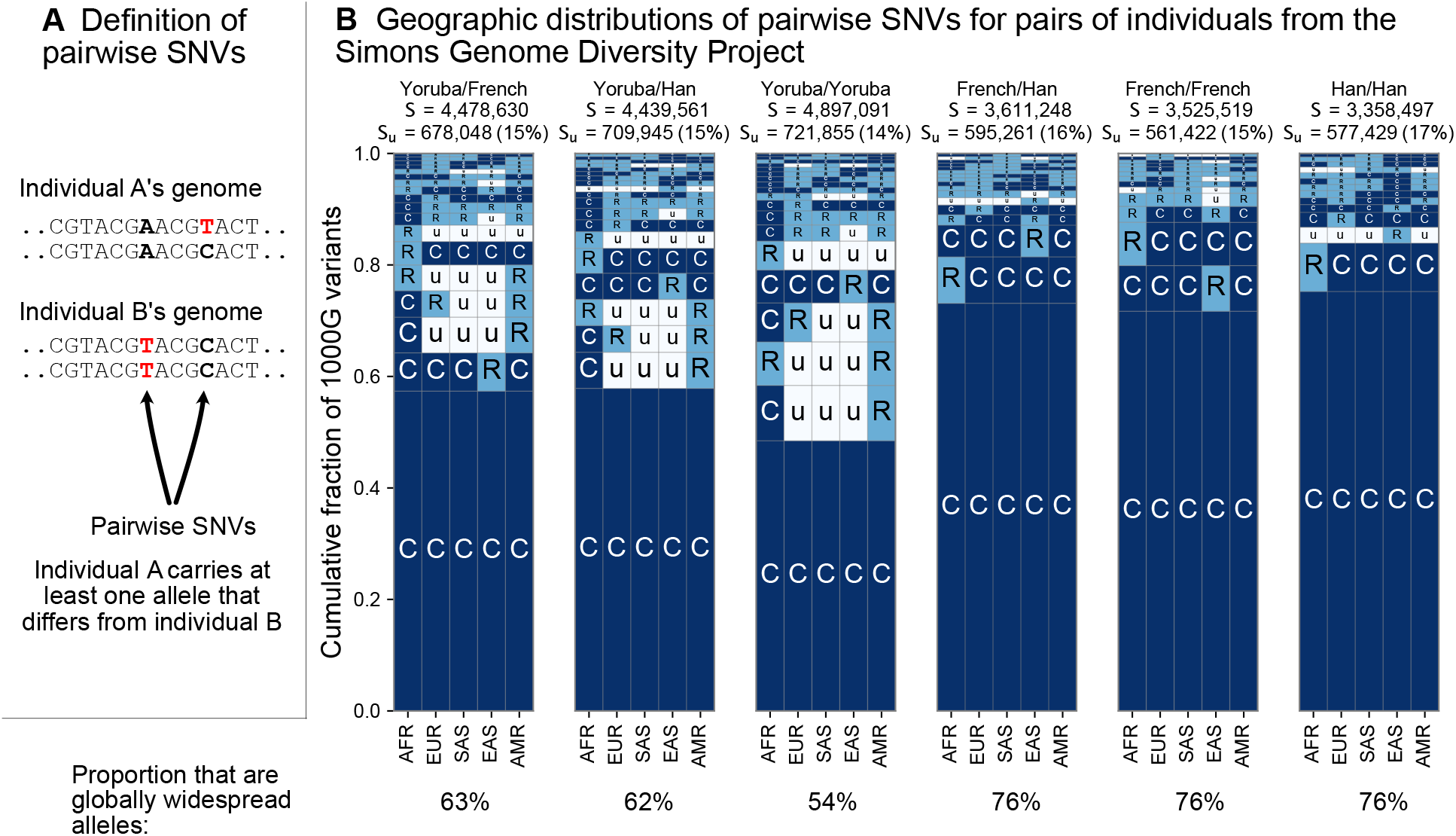
The geographic distributions of SNVs between pairs of individuals. **A:** Definition of a pairwise SNV. **B:** The abundance of geographic distribution codes for different pairs of individuals from the SGDP dataset. Above each plot we show the total number of variants that differ between each individual (*S*) and the number that were unobserved completely in the 1KGP data (*Su*). Across the bottom we show the proportion of variants with globally widespread alleles for each pair. We calculate this as the fraction of variants with no ‘u’ encodings over the total number of variants (*S*). (Note: by doing so, we make the assumption that if a variant is not found in the 1KGP data it is not globally widespread).

From the example pairwise comparisons (Figure 5, and Figure S6), one also observes evidence for higher diversity in Africa, which is typically interpreted in terms of founder effects reducing diversity outside of Africa (Cann, Stoneking, and Wilson 1987; H. C. Harpending and Eller 2000; H. Harpending and Rogers 2000; Ramachandran et al. 2005; Prugnolle, Manica, and Balloux 2005); though other models, especially ones including substantial subsequent admixture, can also produce this pattern (DeGiorgio, Jakobsson, and Rosenberg 2009; Pickrell and Reich 2014). For example, the two Yoruba individuals have more pairwise SNVs (S = 4,897,091) than the French/French (S = 3,525,519) and Han/Han (S = 3,358,497) pairs. Pairs involving one or both of the sample Yoruba individuals have more variants with alleles common in Africa and rare or absent elsewhere (e.g., ‘CuuuR’, ‘RuuuR’). Finally, a more subtle, but expected, impact of founder effects is that the sample Yoruba/Yoruba comparison is expected to have higher numbers of pairwise variants than the sample Yoruba/Han or Yoruba/French comparison, which we observe.

### The geographic distributions of variants typed on genotyping arrays

Targeted genotyping arrays are a cost-effective alternative to whole-genome sequencing. The geographic distribution of the variants on genotyping arrays affects genotype imputation and genetic risk prediction (Howie et al. 2012; Martin et al. 2017). In contrast to whole-genome sequencing, genotyping arrays use targeted probes to measure an individual’s genotype only at preselected variant sites. The process of discovering and selecting these target sites typically enriches the probe sets towards common variants (Clark et al. 2005) and underrepresents geographically localized variants (Albrechtsen, Nielsen, and Nielsen 2010; Lachance and Tishkoff 2013).

Figure 6 shows the geographic distributions of bi-allelic SNVs included on five popular array products in the 1KGP data. In stark contrast with the SNVs identified by whole-genome sequencing (Figure 3B), a large fraction of the variants on genotyping arrays are globally common, especially for the Affy6, Human Origins, and OmniExpress arrays which were designed primarily to capture common variants. The Omni2.5Exome and MEGA arrays in contrast exhibit many more rare variants. In both of these arrays, the second and third most abundant codes are ‘CuuuR’ and ‘RuuuR’ variants. The MEGA array was uniquely designed to capture rare variation in undersampled continental groups, including African ancestries (Bien et al. 2016, 2019). Wojcik et al (2019) found that this design improved African and African American imputation accuracy, leading to greater power to map population-specific disease risk.

**Figure 6.**
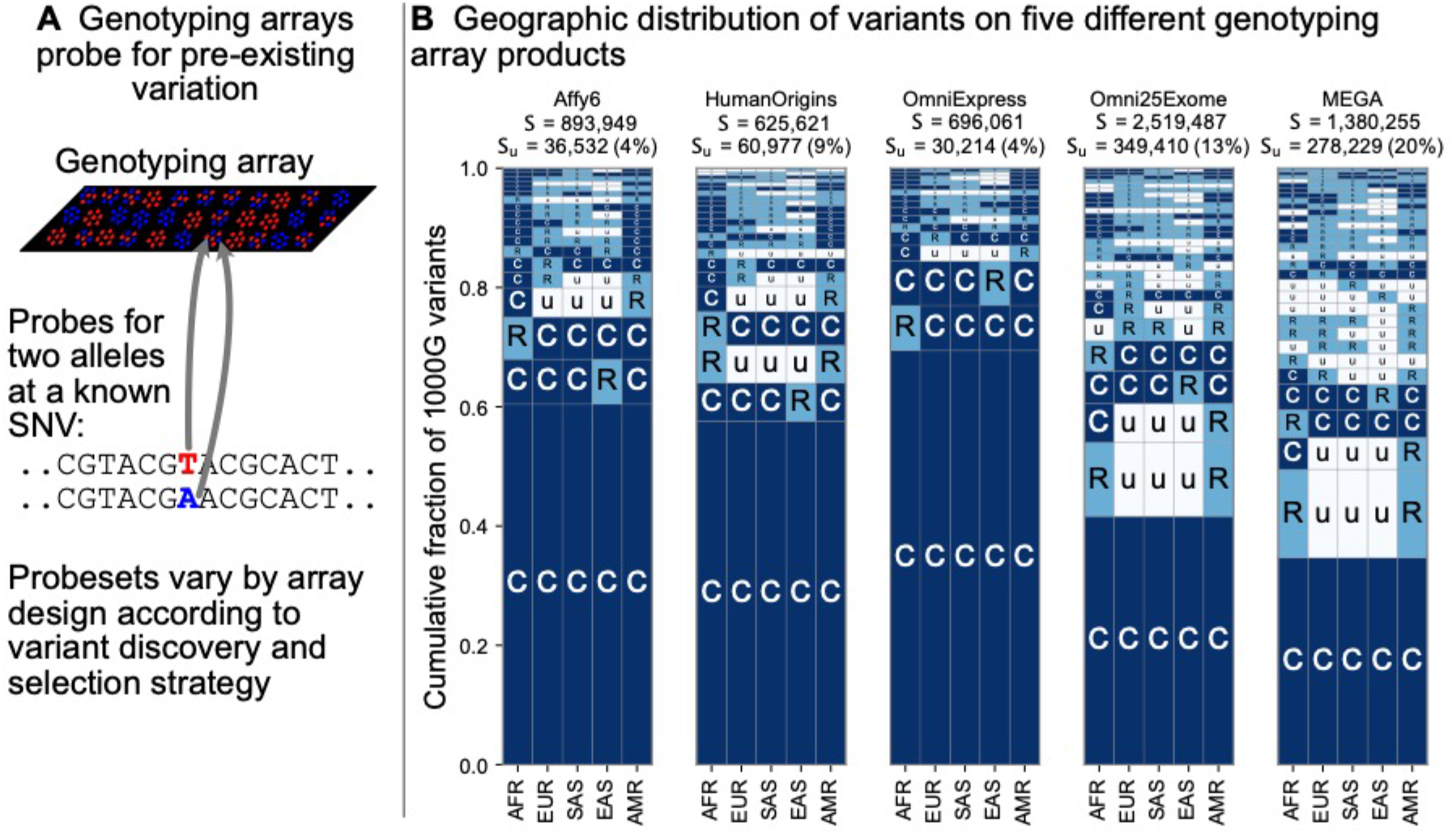
Geographic distribution for variants found on genotyping array products. **A:** Genotyping arrays consist of probes for a fixed set of variants chosen during the design of the array product. **B:** For each array product, we extracted the genomic position of variants found on the array and kept variants that are also found within the 1KGP to highlight their geographic distributions.

### Finer-scale resolution of variant distributions

While the use of 5 regional groupings above allows us to describe variant distributions compactly with a 5-digit encoding, the basic principle of grouping allele frequencies can be extended to build a 26-digit encoding for the 1KGP variants. Doing so, we find a consistent pattern with Figure 2B, in that the majority of variants are seen to be rare and geographically localized (1 ‘R’, and the remainder ‘u’s), and when a variant is common in any one population, it is typically common across the full set of populations (Figure 7, pattern with all ‘C’s). This view reveals that the 5-digit encodings with 1 ‘R’ and 4 ‘u’s are often due to variants that are rare even within a single population. This is not unexpected given many of them are singletons. When we remove singletons (Supp Fig. 7), we again see more clearly rare allele sharing indicative of recent gene flow, though at finer-scale resolution.

**Figure 7.**
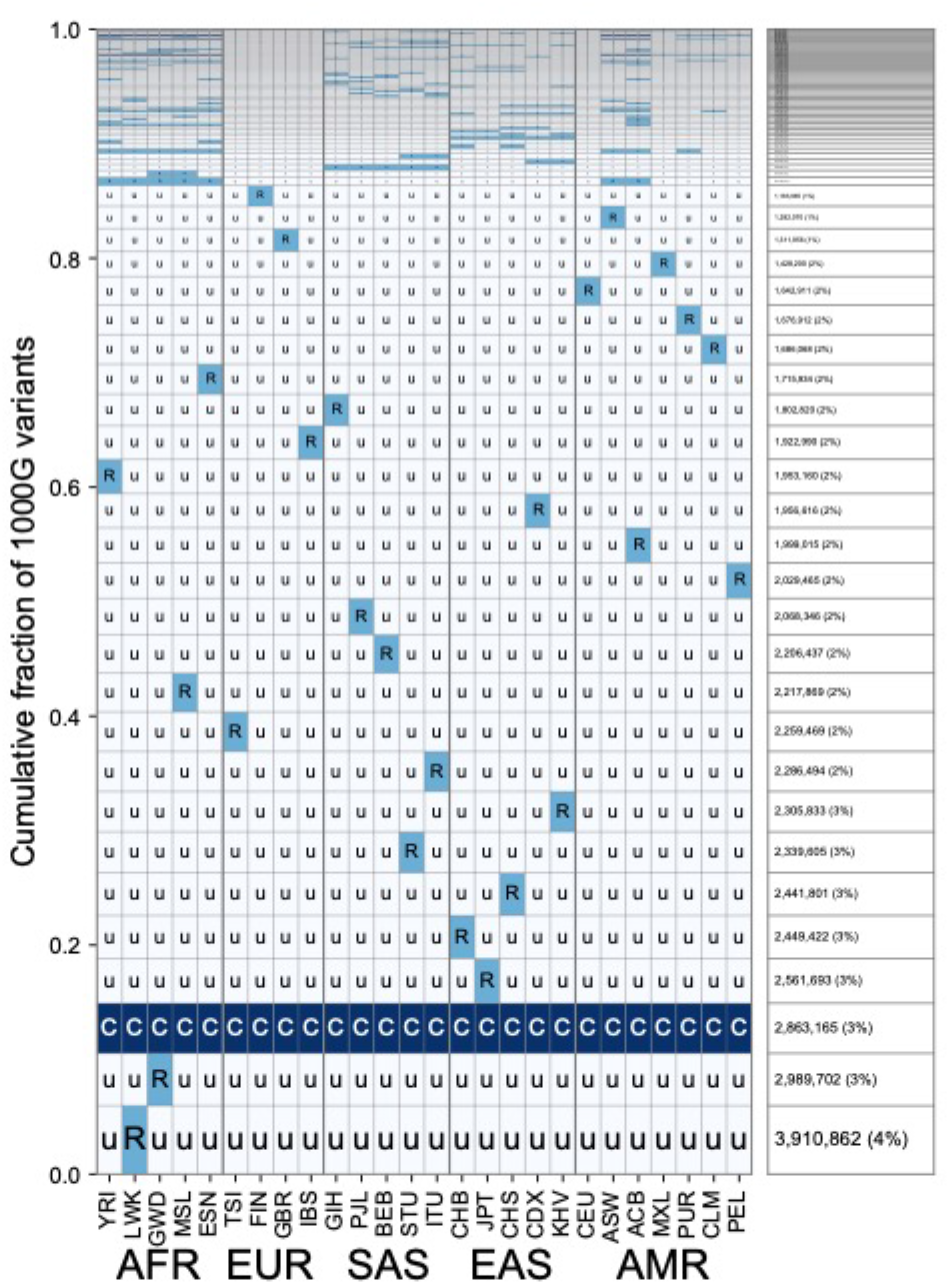
A finer-scale summary of geographic distributions in human SNVs from the 1KGP. This plot is the analogous plot to Figure 3B but rather than calculating frequencies with the 5 regional groupings, we compute them within each of the 26 1KGP populations. The total number of variants represented is the same as in Figure 3B (S = 91,784,367). See Figure 2 for an explanation of the ‘u’,’R’,’C’ codes.

## Discussion

By encoding the geographic distributions of the ~92 million biallelic SNVs in the 1KGP data and tallying their abundances, we have provided a new visualization of human genetic diversity. We term our figures “GeoVar” plots as they help reveal the geographic distribution of sets of variants. GeoVar plots can complement other methods of visualizing population structure, including: plots of pairwise genetic distance, dimensionality-reduction approaches such as PCA, admixture proportion estimates such as STRUCTURE, and explicitly spatial methods that use the sampling locations of individuals (Guillot et al. 2009; Novembre and Peter 2016; Bradburd and Ralph 2019). These previously developed methods help reveal population structure, infer genetic ancestry, and measure historical migration patterns. However, they do a poor job of showing how alleles are distributed geographically. To minimize confusion about levels of differentiation among populations, researchers and educators can consider complementing PCA or STRUCTURE outputs with a variant-centric visualization like the ones presented here. To that end, we provide source code to replicate our figures and to generate similar plots for other datasets (the “GeoVar” software package; see URLs).

A goal of our work was to build a visualization that can help correct common misconceptions about human genetic variation. First, because many existing methods to describe population structure emphasize between-group or between-individual differentiation, they can convey a misleading impression of “deep” divergence between populations when it may not exist. Comparing Figure 1 to outputs of models with “deep” or “shallow” divergence can help teach how patterns of human variation are consistent with shallow divergence and the Recent African Origins model (Box 2). Second, because personal ancestry tests can identify ancestry to broad continental regions, it is possible to incorrectly conclude human alleles are typically found exclusively in a single region and at high frequency within that region (e.g., patterns such as ‘uuCuu’.) As our figures show, this is not the case. Rather, it should be kept in mind that most fine-scale personal ancestry tests work using genotyping arrays and combining evidence from subtle fluctuations in the allele frequencies of many common variants (Novembre and Peter 2016). Finally, another related misconception is that two humans from different regions of the world differ mainly due to alleles that are typical of each region. As we show in Figure 5, most of the variants that differ between two individuals are variants with alleles that are globally widespread.

Our method requires computing allele frequencies within predefined groupings. Grouping and labeling strategies vary between genetic studies and are determined by the goals and constraints of a particular study (Race, Ethnicity, and Genetics Working Group 2005; Panofsky and Bliss 2017; Mathieson and Scally 2020). While we chose deliberately coarse grouping schemes to address the misconceptions described above, the key facts we derive about human genetic variation are robust and appear in finer-grained 26-population versions of the plot (Figure 7). We recommend that any application of the GeoVar approach needs to be interpreted with the choice of groupings in mind.

The visualization method developed here is also useful for comparing the geographic distributions of different subsets of variants, (e.g., Figures 5 and 6). For example, when applied to the list of variants targeted by a genotyping array (Figure 6), the approach quickly reveals the relative balance of common versus rare variants and the geographical patterns of those variants.

Interpreting the results of this visualization approach does have some caveats. First, we estimate the frequency of alleles from samples of local populations. We expect that as sample sizes increase many alleles called as unobserved ‘u’ will be reclassified as rare ‘R’. The average sample size across all of our geographic regions is approximately 500 individuals (AFR: 504, EUR: 404, SAS: 489, EAS: 504, AMR: 603). Assuming regions are internally well-mixed, we have ~80% power to detect alleles with a frequency of ~0.2% in a region (Figure S4). For alleles with lower frequencies, we would require larger sample sizes to ensure similar detection power. An implication is that in large samples, we should observe more rare variant sharing. Thus, we expect the figures here to underrepresent the levels of rare variant sharing between human populations.

A second caveat is that our encoding groups a wide range of variants into the “(C)ommon” category (i.e., all variants where the frequency of the globally minor allele is greater than 5%). For some applications, such as population screening for carriers, it may be enough to know that a variant falls in the “rare” or “common” bins we have described, and more detail is inconsequential. For other applications, the detailed fluctuations in allele frequency across populations are relevant—for example, differences in allele frequencies at common variants (Figure S5) are regularly used to infer patterns of population structure and relatedness (Li et al. 2008; Pickrell and Pritchard 2012; Patterson et al. 2012).

Third, one must interpret our results with the sampling design of the 1KGP study design in mind. In particular, the 1KGP filtered for individuals of a single ethnicity within each locale. However, in our current cosmopolitan world, the genetic diversity in any location or broad-based sampling project will be considerably higher than implied by the geographic groupings above. For example, the UK Biobank, while predominantly of European ancestry, has representation of individuals from each of the five regions used here (Bycroft et al. 2018). The 1KGP also sampled South Asian ancestry from multiple locations outside of South Asia, and whether those individuals show excess allele sharing due to recent admixture in those contexts is unclear. While we expect overall similar patterns to those seen here using emerging alternative datasets (Bergström et al. 2019), there may be subtle differences due to sampling and study design considerations.

Despite these caveats, the results of the visualizations provided here help reinforce the conclusions of a long history of empirical studies in human genetics (Lewontin 1972; Ramachandran et al. 2005; Conrad et al. 2006; Li et al. 2008; 1000 Genomes Project Consortium et al. 2015; Mallick et al. 2016; Bergström et al. 2019). The results show how the human population has an abundance of localized rare variants and broadly shared common variants, with a paucity of private, locally common variants. Together these are footprints of the recent common ancestry of all human groups. As a consequence, human individuals most often differ from one another due to common variants that are found across the globe. Finally, though not examined explicitly above, the large abundance of rare variants observed here is another key feature of human variation and a consequence of recent human population growth (Slatkin and Hudson 1991; Di Rienzo and Wilson 1991; Keinan and Clark 2012; Nelson et al. 2012; Tennessen et al. 2012).

The well-established introgression of archaic hominids (e.g., Neandertals, Denisovans) into modern human populations (Wolf and Akey 2018) is not apparent in the GeoVar plots we produced. We believe that there are two broad reasons for this: (1) The clearest signal of archaic introgression will come from sites where archaic hominids differed from modern humans, and we expect that these sites are only a very small fraction of variants found in humans today. The average human–Neandertal and human–Denisovan sequence divergence are both less than 0.16% (using observations from Prüfer et al. 2014), and a recent study estimates that there are fewer than 70 Mb (2.3% of the genome) of Neanderthal introgressed segments per individual for all individuals in the 1KGP (Chen et al. 2020). (2) We do not expect SNVs from archaic introgression to be concentrated in a single GeoVar category. For example, introgressed variants occupy a wide range of allele frequencies (Bergström et al. 2019). Archaic introgression events are believed to be old: >30,000 years ago, allowing time for substantial genetic drift and admixture among human populations (Chen et al. 2020). Negative selection (Harris and Nielsen 2016; Juric, Aeschbacher, and Coop 2016) and, in some cases, strong positive selection (Racimo et al. 2015) have also shaped the patterns of introgressed SNVs. For these reasons, we expect low levels of archaic introgression not to create a striking visual deviation in our GeoVar plots from the background patterns of a Recent African Origin model with subsequent migration (Box 2). To highlight the contributions of archaic hominids to human variation, more targeted approaches are needed (e.g., Green et al. 2010; Durand et al. 2011). Future work could also naturally extend the approach here to include archaic sequence data.

The geographic distributions of genetic variants visualized here are relevant for a number of applications, including studying geographically varying selection (Yi et al. 2010; Key et al. 2018), human demographic history (Gutenkunst et al. 2009), and the genetics of disease risk. For instance, due to ascertainment bias in arrays (Figure 6) and power considerations, common variants are often found in genome-wide association studies of disease traits (Manolio et al. 2009). The patterns shown above make it clear that most common variants are shared across geographic regions. Indeed, many common variant associations replicate across populations (Marigorta and Navarro 2013; though see Martin et al. 2017; Mostafavi et al. 2020 for complications). More recently, due to increasing sample sizes and sequencing-based approaches, disease mapping studies are finding more associations with rare variants (Bomba, Walter, and Soranzo 2017). As our work here emphasizes, rare variants are likely to be geographically restricted, and so one can expect the rare variants found in one population will not be useful for explaining trait variation in other populations, though they may identify relevant biological pathways that are shared across populations.

A future direction for the work here would be to apply our approach to other classes of genetic variants such as insertions, deletions, microsatellites, and structural variants. We note that in studies with sample sizes similar to or smaller than the 1KGP, nearly all SNVs arise from single mutation events. For other variants that arise from single mutation events (e.g., indels that arise from single mutations), we expect similar patterns to those observed for SNVs here. In contrast, for highly mutable loci, such as microsatellites, we expect alleles will be distributed in disjoint regions of the world due to multiple mutational origins (Ralph and Coop 2013; Mathieson and McVean 2014; Phillips et al. 2020).

Another future direction would be to shift from visualizing patterns of allele sharing to the patterns of sharing of ancestral lineages in coalescent genealogies. Recent advances in the inference of genome-wide tree sequences (Kelleher et al. 2019; Speidel et al. 2019) and allele ages (Albers and McVean 2020) allow for quantitative summaries of ancestral lineage sharing. Such quantities have a close relationship to the multi-population SFS properties that are studied here, yet are more fundamental in a sense and less subject to the stochasticity of the mutation process. That said, the conceptual simplicity of visualizing allele frequency patterns may be an advantage in educational settings.

Most importantly, future applications of the approach will ideally use datasets that include a greater sampling of the world’s genetic diversity (Bustamante, Burchard, and De la Vega 2011; Popejoy and Fullerton 2016; Martin et al. 2017; Peterson et al. 2019). A related point is that the application of our method to genotyping array variants (Fig. 6) reinforces the importance of considering the ancestry of study populations in genotype array design and selection (Peterson et al. 2019).

Overall, the visualizations produced here provide an interpretable way to depict geographic patterns of human genetic variation. With personal genomic technologies and ancestry testing becoming commonplace, there is increasing importance in fostering the understanding of human population genetics. To this end, human genetics researchers must develop interpretable materials on patterns of genetic variation for use in educational and outreach settings (Donovan et al. 2019). The variant-centric approach detailed here complements existing visualizations of population structure, facilitating a clearer understanding of the major patterns of human genetic diversity.

## Acknowledgments

The 1000 Genomes data used here were generated at the New York Genome Center with funds provided by NHGRI Grant 3UM1HG008901-03S1. We thank members of the Novembre Lab, especially as this project was initiated in a group hackathon with contributions from Hussein Al-Asadi, Kushal Dey, Evan Koch, Joe Marcus, Ben Peter, Mark Reppell, and Joel Smith. We also thank Jeremy Berg, Jedidiah Carlson, Anna Di Rienzo, Joe Marcus, Aaron Panofsky, Molly Przeworski, Harald Ringbauer, Mashaal Sohail, Matthias Steinrücken, and Xin He for comments on the manuscript draft, and Paul Strode and Brian Donovan for helpful conversations. This work was completed in part with resources provided by the University of Chicago’s Research Computing Center and was supported by NIH training grant T32 GM07197 (AB), the University of Chicago “Chicago Fellows” program (DPR), and NIH grant R01 GM132383.

## URLS

1. *1000 Genomes High-Coverage Data:* http://ftp.1000genomes.ebi.ac.uk/vol1/ftp/data_collections/1000G_2504_high_coverage/working/20190425_NYGC_GATK/
2. *GCRh38 Genome Masks (describing the “accessible” genome):* http://ftp.1000genomes.ebi.ac.uk/vol1/ftp/data_collections/1000_genomes_project/working/20160622_genome_mask_GRCh38/
3. *Simons Genome Diversity Project Data:* https://reichdata.hms.harvard.edu/pub/datasets/sgdp/
4. *Genotyping Array Variant Lists:*

a. *Human Origins Array:* https://sec-assets.thermofisher.com/TFS-Assets/LSG/Support-Files/Axiom_GW_HuOrigin.na35.annot.csv.zip
b. *Affymetrix GenomeWide 6.0 Array:* http://www.affymetrix.com/Auth/analysis/downloads/na35/genotyping/GenomeWideSNP_6.na35.annot.csv.zip
c. *Illumina MEGA Array:* ftp://webdata2:webdata2@ussd-ftp.illumina.com/downloads/productfiles/multiethnic-amr-afr-8/v1-0/multi-ethnic-amr-afr-8-v1-0-a1-manifest-file-csv.zip
d. *Illumina Human Omni Express:* ftp://webdata:webdata@ussd-ftp.illumina.com/Downloads/ProductFiles/HumanOmniExpress-24/v1-0/HumanOmniExpress-24-v1-0-B.csv
e. *Illumina Omni2.5Exome:* ftp://webdata:webdata@ussd-ftp.illumina.com/Downloads/ProductFiles/HumanOmni2-5Exome-8/Product_Files_v1-1/HumanOmni2-5Exome-8-v1-1-A.csv
5. *Ancestral allele calls:* ftp://ftp.ensembl.org/pub/release-90/fasta/ancestral_alleles/homo_sapiens_ancestor_GRCh38_e86.tar.gz
6. *Pipeline for reproducing results:* https://github.com/aabiddanda/geovar_rep_paper

## Supplementary Tables and Figures

**Supplementary Table 1.** Table of abbreviations and groupings used in the study. The regional groupings used in this study only differ in having CEU, ASW and ACB assigned to the AMR regional group.

| **Population description** | **Population code (1KGP)** | **Superpopulation code (1KGP)** | **Regional grouping (This study)** |
| --- | --- | --- | --- |
| African Ancestry in Southwest US | ASW | AFR | AMR |
| Yoruba in Ibadan, Nigeria | YRI | AFR | AFR |
| Luhya in Webuye, Kenya | LWK | AFR | AFR |
| Japanese in Tokyo, Japan | JPT | EAS | EAS |
| Chinese Dai in Xishuangbanna, China | CDX | EAS | EAS |
| Utah residents (CEPH) with Northern and Western European ancestry | CEU | EUR | AMR |
| Han Chinese in Beijing, China | CHB | EAS | EAS |
| Gujarati Indians in Houston, TX | GIH | SAS | SAS |
| Gambian in Western Division, The Gambia - Mandinka | GWD | AFR | AFR |
| Mende in Sierra Leone | MSL | AFR | AFR |
| Iberian populations in Spain | IBS | EUR | EUR |
| Toscani in Italy | TSI | EUR | EUR |
| Punjabi in Lahore, Pakistan | PJL | SAS | SAS |
| Esan in Nigeria | ESN | AFR | AFR |
| Mexican Ancestry in Los Angeles, California | MXL | AMR | AMR |
| Colombian in Medellin, Colombia | CLM | AMR | AMR |
| Peruvian in Lima, Peru | PEL | AMR | AMR |
| Han Chinese South | CHS | EAS | CHB |
| Puerto Rican in Puerto Rico | PUR | AMR | AMR |
| Finnish in Finland | FIN | EUR | EUR |
| Kinh in Ho Chi Minh City, Vietnam | KHV | EAS | EAS |
| Bengali in Bangladesh | BEB | SAS | SAS |
| African Caribbean in Barbados | ACB | AFR | AMR |
| Indian Telugu in the UK | ITU | SAS | SAS |
| British in England and Scotland | GBR | EUR | EUR |
| Sri Lankan Tamil in the UK | STU | SAS | SAS |

**Supplementary Figure 1.**
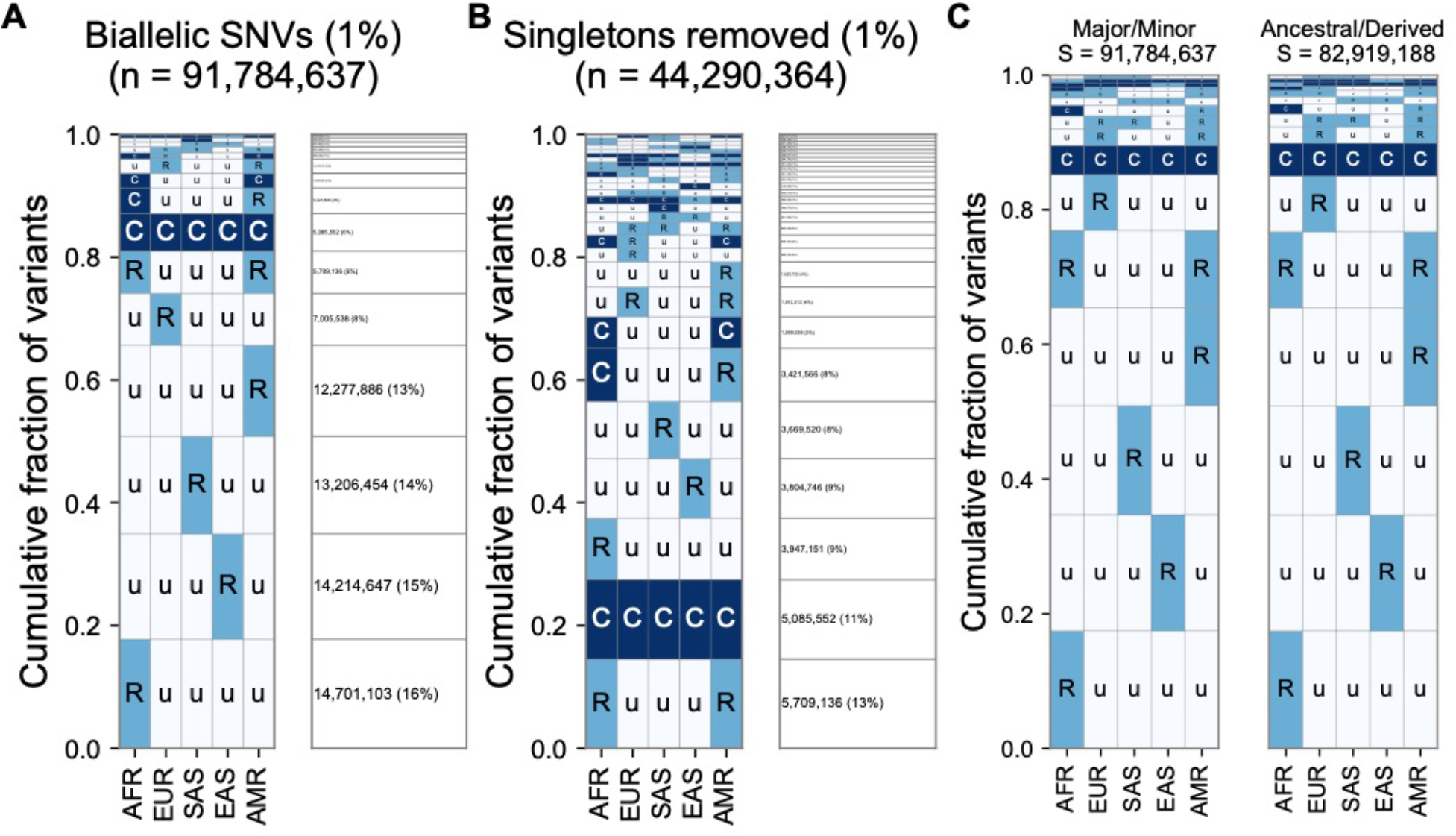
**A:** The relative abundance of geographic distribution codes within the ~92 million variants when using an MAF of 1% as the distinction between “common” (‘C’), and “rare” (‘R’). The right-hand panel shows the percentage of variants that fall within the geographic code represented on the left-hand side; distribution patterns are sorted by their abundance, from bottom-to-top. **B:** The abundance of geographic distribution codes for ~44 million nonsingleton variants using an MAF of 1% as the boundary between “common” (‘C’), and “rare” (‘R’). **C:** Comparison for the abundance of geographic distribution codes when polarizing to the ancestral and derived allele (using build 38) versus major/minor allele. We only include positions where an ancestral allele is supported by at least two outgroups. At 96.6% of variants (80,068,013 / 82,919,198), the minor allele is also the derived allele.

**Supplementary Figure 2.**
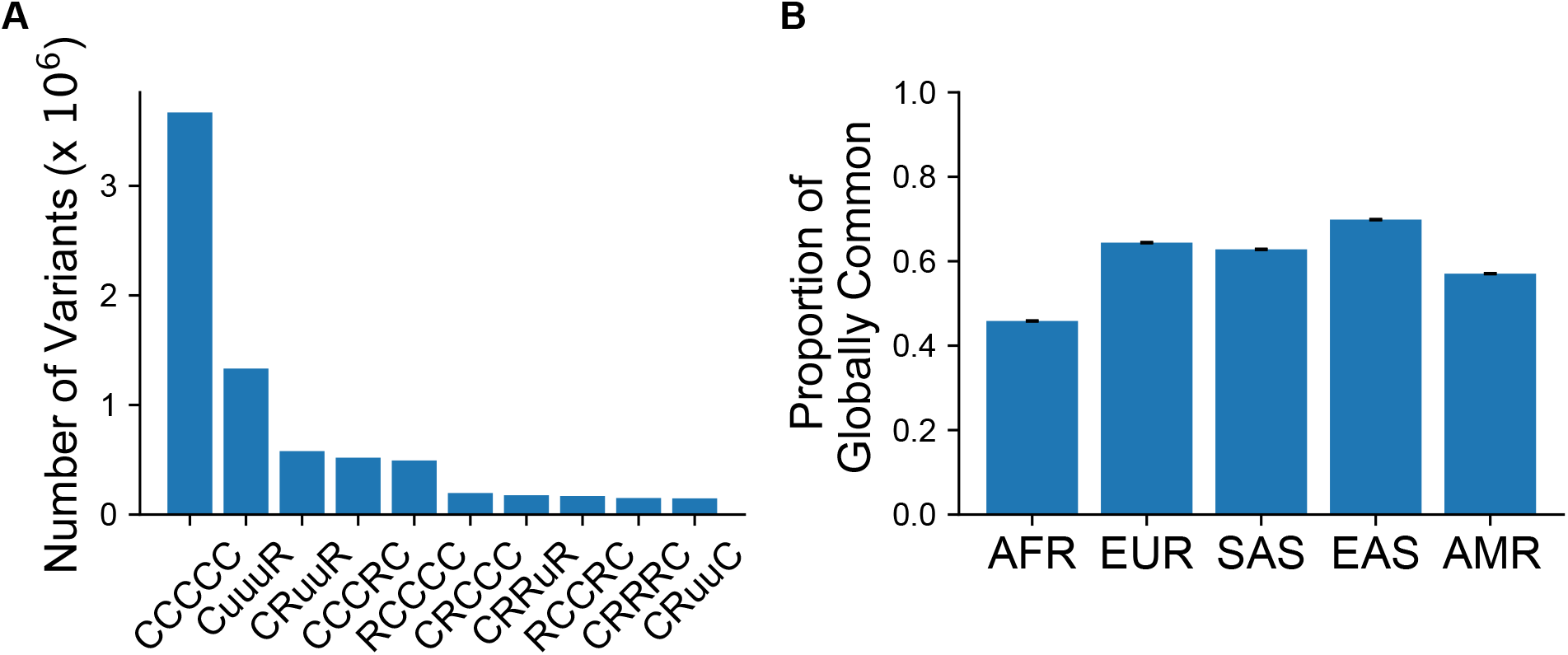
Top 10 categories when conditioning on the variant being “common” (MAF > 5%) in at least one population. Conditioned on a variant being common in a single region, 37.3% of variants are categorized as “globally common” or “CCCCC”. **B:** The proportion of variants that fall within the “globally common” or “CCCCC” geographic distribution code conditional on the variant being common (MAF > 5%) in the specific continental group

**Supplementary Figure 3.**
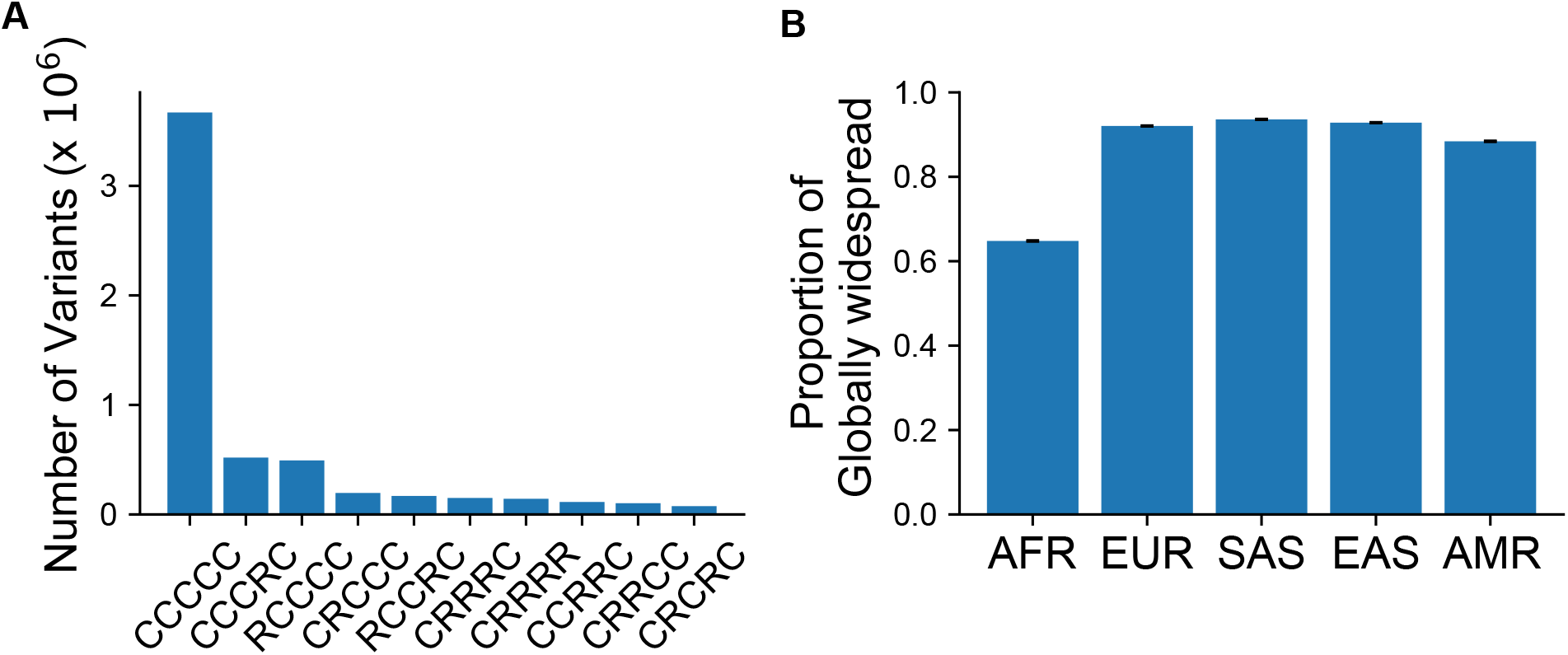
**A:** The number of variants that fall within a given geographic distribution code conditional on the variant being “globally widespread”, i.e. a category that has no unobserved (“u”) codes. We note that 55.6 % of variants conditioned on being globally widespread are also globally common (“CCCCC”). In terms of absolute numbers, variants that are common in at least one population (S = 9,958,838) that are also globally widespread (S = 6,322,767) comprise ~ 63% of the total when conditioning on being common in at least one population. When conditioning on variants common only in regions outside Africa (S = 7,544,648), the percentage of globally widespread variants (S = 6,179,781) increases to ~ 82 %. **B:** The proportion of variants that fall within a “globally present” category, defined as categories that contain no unobserved (“u”) codes, conditional on the variant being common (MAF > 5%) in the specific continental group

**Supplementary Figure 4.**
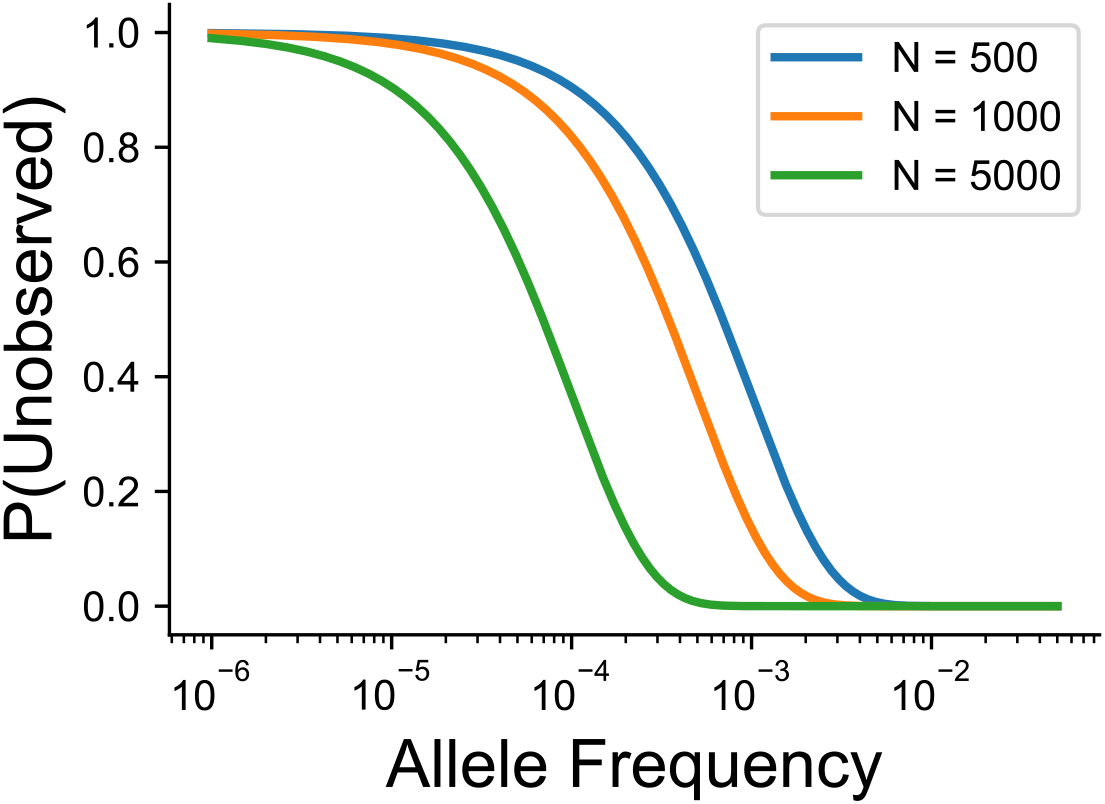
Probability of not observing a variant at a given allele frequency and sample size in number of individuals. We have assumed that allele frequencies follow Hardy-Weinberg equilibrium, and the probability of no observations of an allele is calculated using the binomial distribution.

**Supplementary Figure 5.**
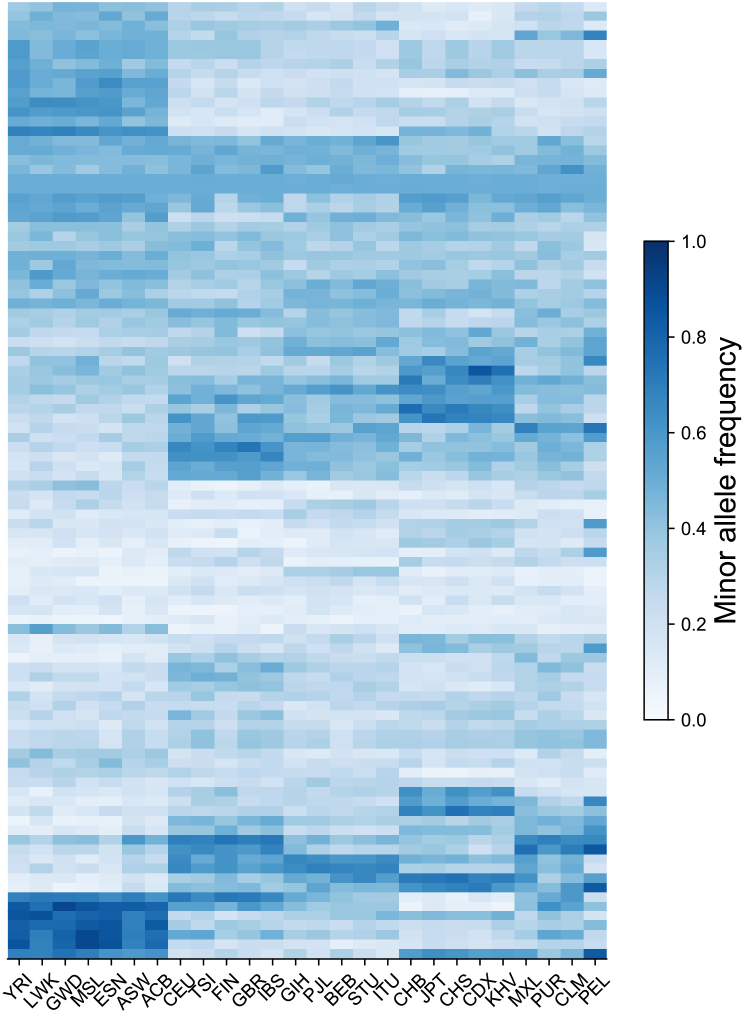
Heatmap of MAF across all 26 original population labels in the 1KGP using 300 randomly chosen variants on Chromosome 22 that have MAF > 5% in all populations. Variants are ordered based on hierarchical clustering on the Euclidean distance between minor allele frequency profiles.

**Supplementary Figure 6:**
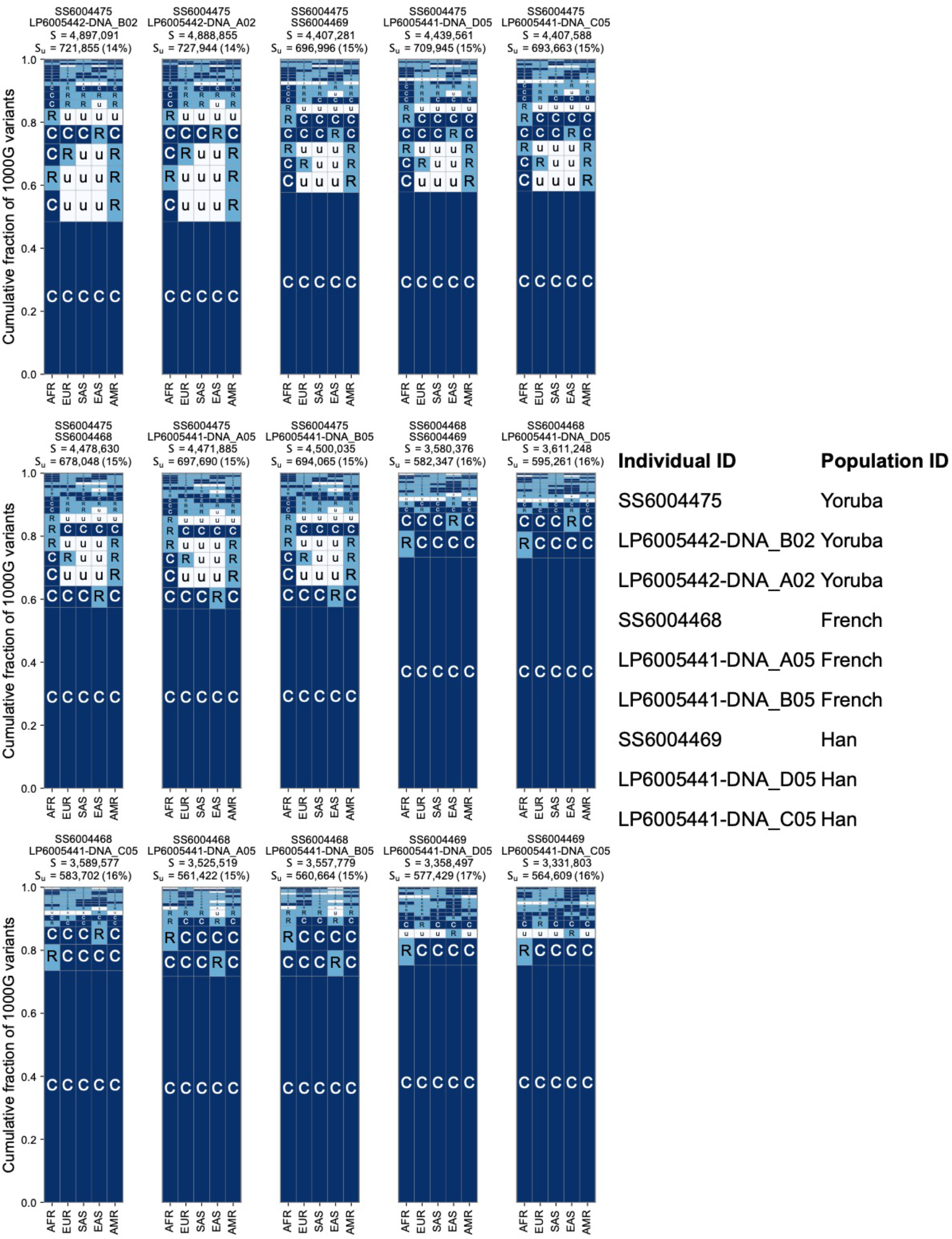
Additional examples of geographic distribution codes for pairwise variants from pairs of sampled individuals in the SGDP.

**Supplementary Figure 7.**
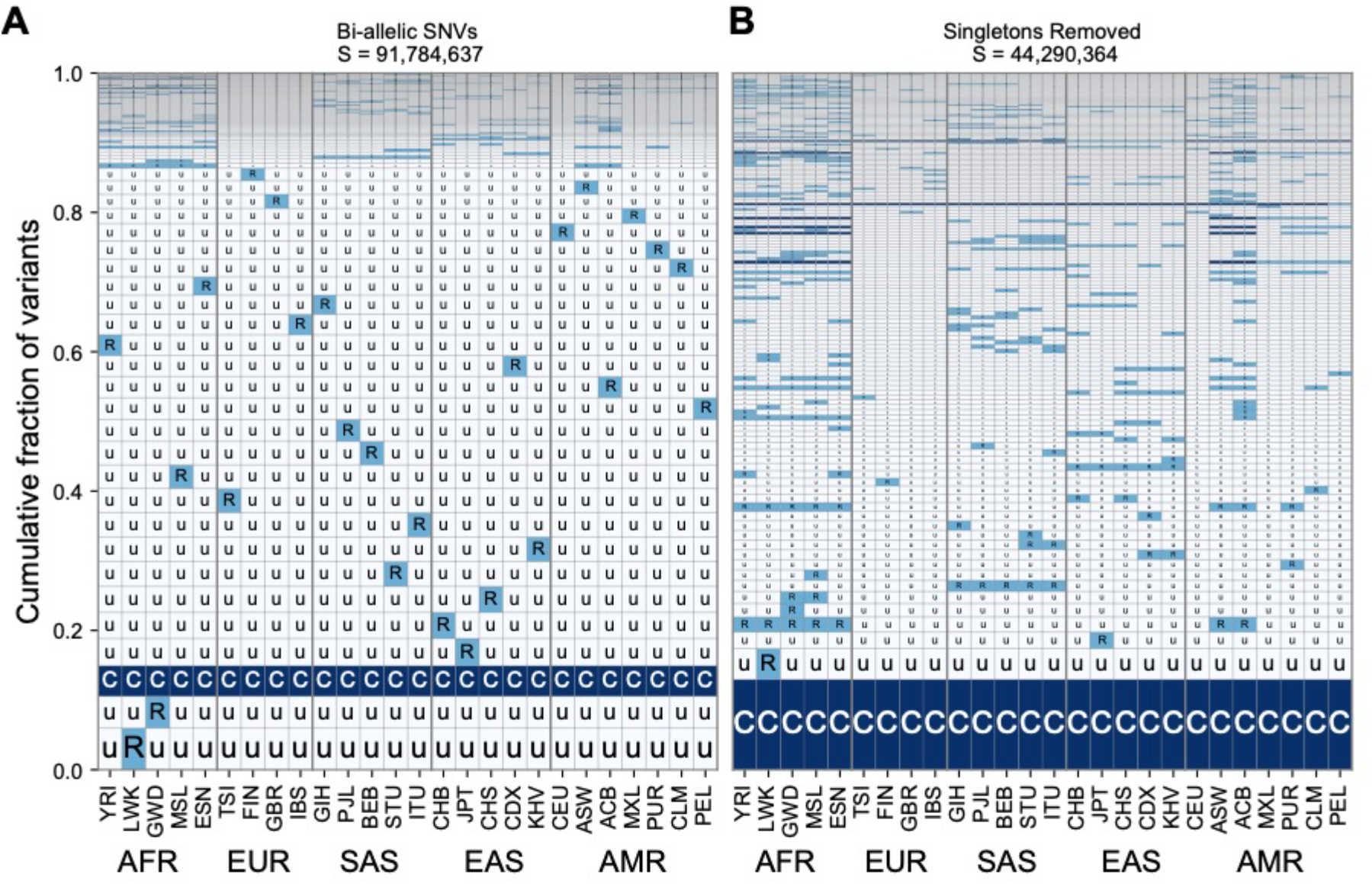
The geographic distribution of variants across all 26 populations (Table S1) in the 1KGP both with singletons included (A) and removed (B). Regional groupings are provided on the bottom to reflect our choices for population groupings throughout the main paper.

**Supplementary Figure 8.**
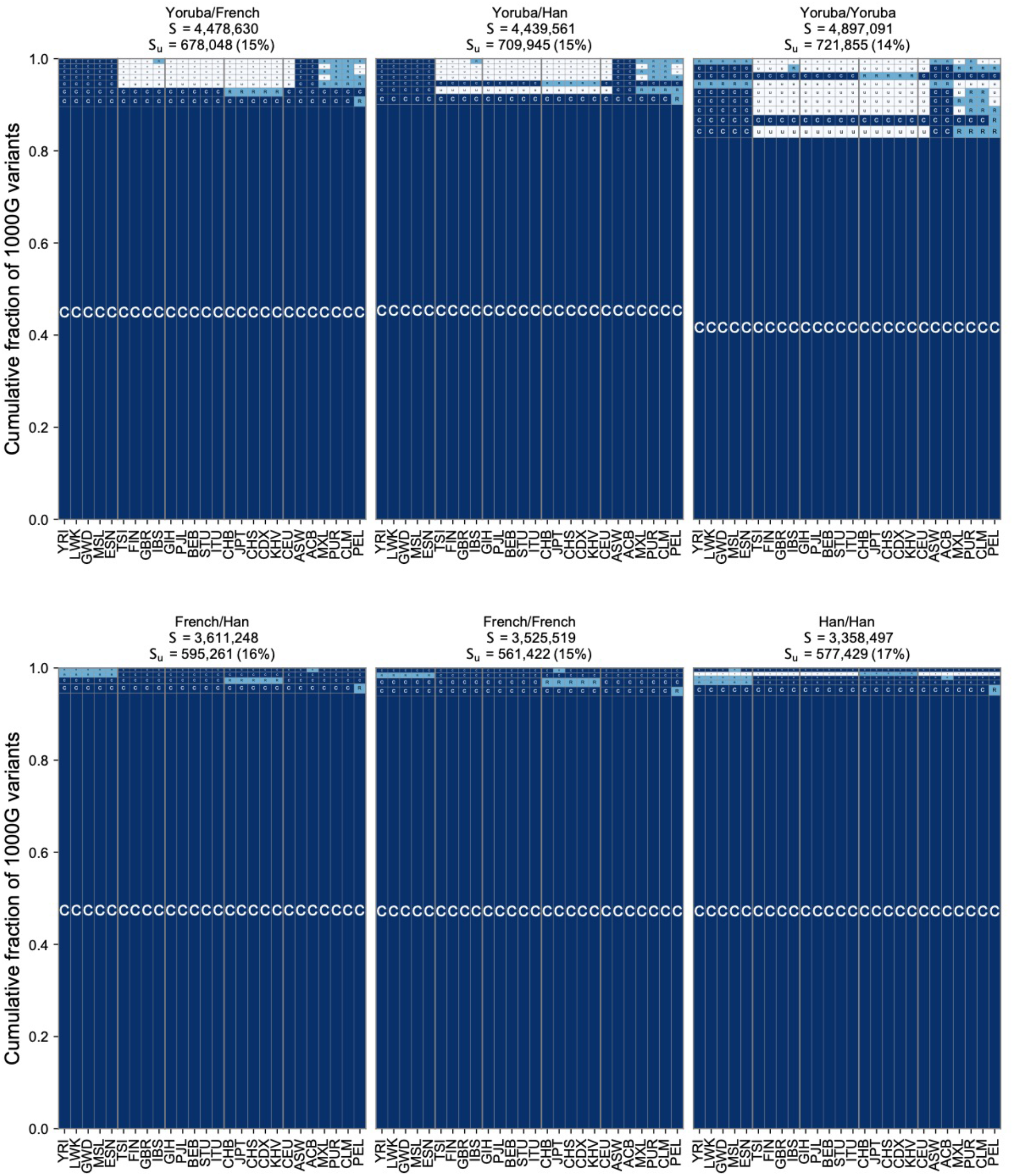
The geographic distribution of pairwise SNVs across pairs of individuals from the Simons Genome Diversity Project using the full set of 26 populations from the 1KGP.

**Supplementary figure 9.**
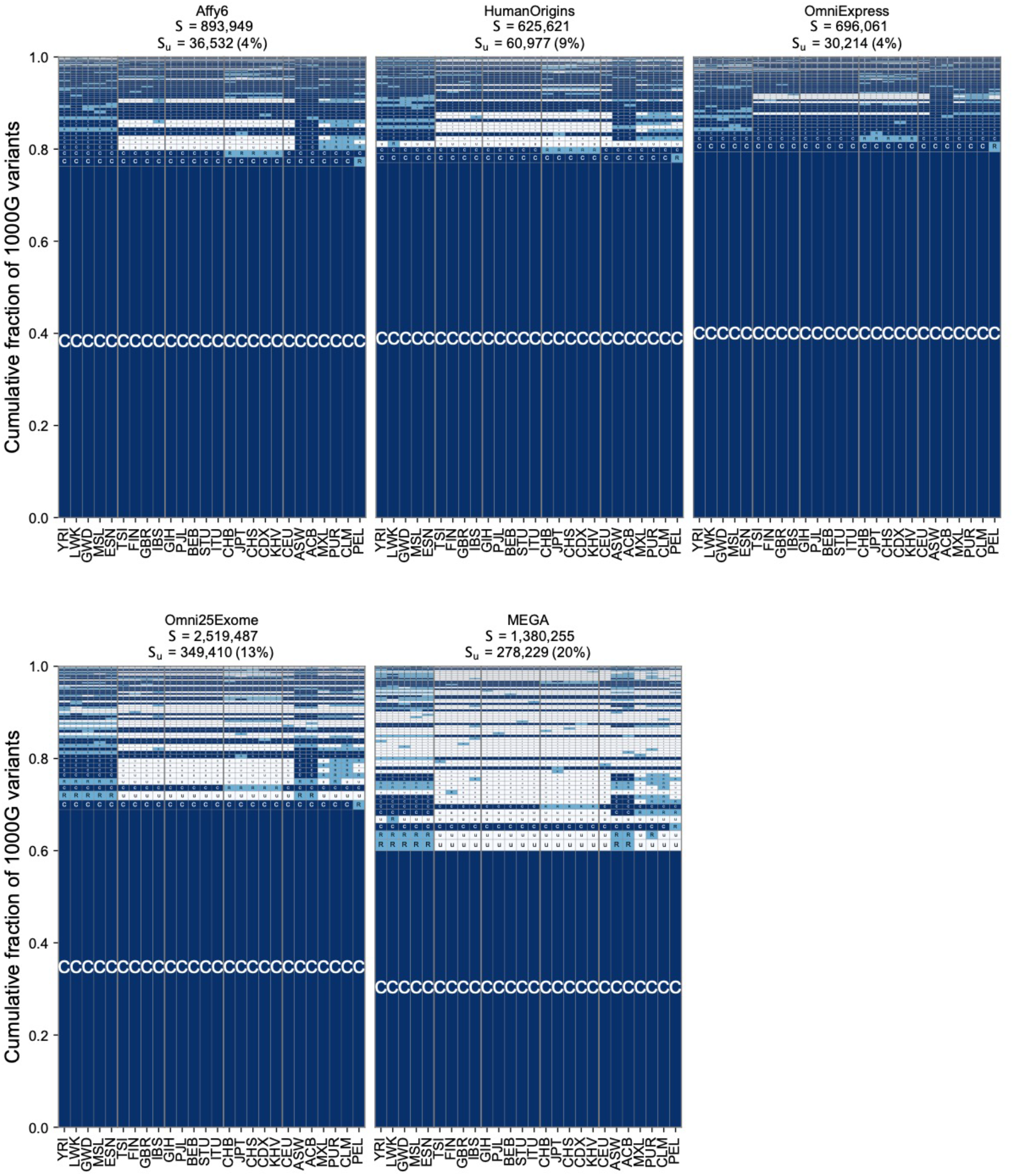
The geographic distribution of SNVs on genotyping arrays using the full set of 26 populations from the 1KGP.

## Appendix A: Theoretical geographic distribution code abundances

The relative abundances of geographic distribution codes derive from human population history (Box 2). Here, we use a simple population genetic model to develop intuition about the relationship between the divergence time of a pair of populations and the expected two-letter code abundances. To isolate the effect of population divergence from other factors such as population growth, we consider the simplest possible model of divergence: two constant-size populations of N individuals descended from a single N-individual source population *T* generations ago (Fig. 4A). We incorporate recent contact between populations via a symmetric admixture coefficient *α*. Individuals in Population 1 derive a fraction *α* of their ancestry from Population 2 and vice versa. Human population history is much more complex than our model, but it captures the essential features of common ancestry, subsequent isolation, and modern admixture.

Python source code implementing the calculation and producing Fig. 4 is available in the project’s Git repository (https://github.com/aabiddanda/geovar_rep_paper).

### Wright-Fisher diffusion of allele frequencies

In our model, allele frequencies in the two source populations are initially identical because they derive from the same source population. After the populations split, allele frequencies evolve independently according to a Wright-Fisher diffusion with symmetric mutations at rate *θ* new mutations per population per generation. At time *t = T*/2*N* generations after the split, the joint density of mutations at frequency *x*_1_ in Population 1 and x_2_ in Population 2 is given by,

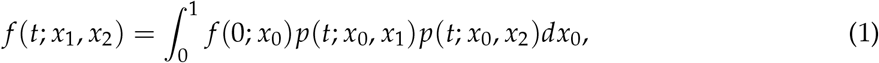

where *f* (0; *x*_0_) is the density of mutations at frequency *x*_0_ in the source population and *p*(*t*; ·, ·) is the Wright-Fisher transition density function. Assuming that the source population was at mutation-drift equilibrium, *f* (0; *x*_0_) = *π*(*x*_0_) ∝ (*x*_0_(1 − *x*_0_))^*θ*−1^, the stationary measure of the Wright-Fisher diffusion.

We use the spectral decomposition of Song and Steinrücken [1] to represent the Wright-Fisher transition density as an infinite sum of modified Jacobi polynomials, *B_i_*(*x*):

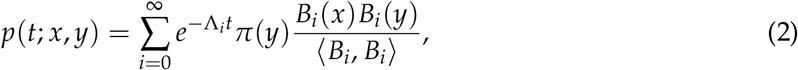

where the inner product 〈*g, h*〉 is given by 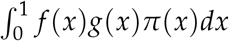 The Jacobi polynomials are orthogonal with respect to this inner product. That is, 〈*B_i_, B_j_*〉 = 0 for *i ≠ j*. Substituting (2) into (1) and using orthogonality, we have:

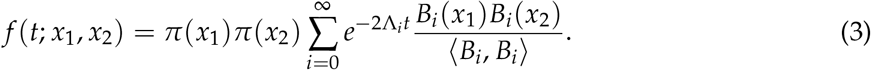

In practice, we can only compute partial sums on the right-hand side, which we can re-write as

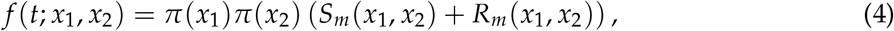

where *S_m_* is the partial sum of terms up to order *m* and *R_m_* is the remainder, which represents the error from truncating the series. We can control this error by choosing a large enough *m* (see Numerical Integration.)

### Sampling probabilities

The abundances of two-population distribution codes is a simple transformation of the cumulative distribution function (CDF) of the joint allele counts (*K*_1_, *K*_2_). Conditioning on allele frequencies at time t, but before admixture, the CDF is given by

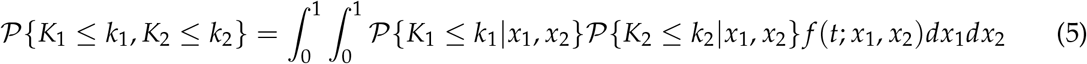

For *n* randomly sampled haploid individuals from each population, and admixture coefficient *α*, we have:

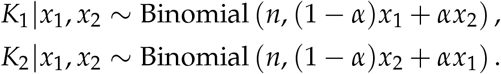

Writing 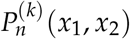 for the binomial cumulative distribution function 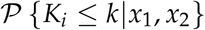, and substituting (5) into (4) yields:

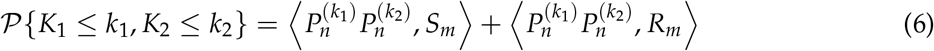

where the inner product now represents the double integral weighted by *π*(*x*_1_)*π*(*x*_2_).

### Numerical integration

We compute the integrals in (6) by two-dimensional Gauss-Jacobi quadrature. The left argument of the inner product is a polynomial of degree *n* in both *x*_1_ and *x*_2_. As a result, we can choose *m* = 2*n*, so that 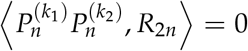 due to the orthogonality of the Jacobi polynomials. Because *S*2_*n*_ is also a polynomial, the integrand is a polynomial of degree 4*n*. Thus, fixed-order tensor-product Gauss-Jacobi quadrature is guaranteed to yield the exact integral with 4*n*^2^ evaluations of the integrand.

## Appendix B: Extinction probability and conditional mean frequency

The extinction probability 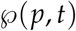, the probability that a mutation that was at frequency *p* at time *t* = 0 is extinct at time *t = T*/2*N*, obeys the Kolmogorov backward equation [2]:

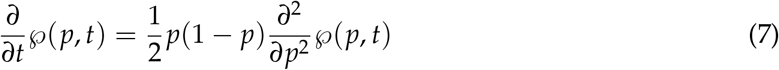

with boundary conditions

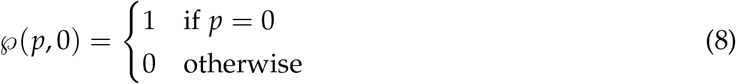

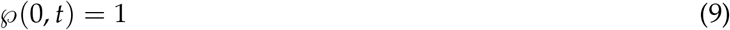

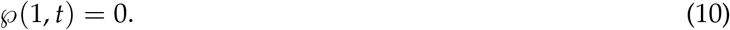

For short times and rare alleles (i.e., *t, p* ≪ 1), we can use the approximation *p*(1 − *p*) ≈ *p*, to get a simpler diffusion equation:

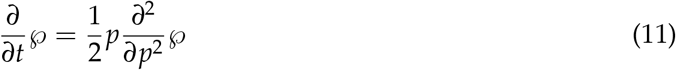

We can solve this equation in closed form to find the time-dependent extinction probability,

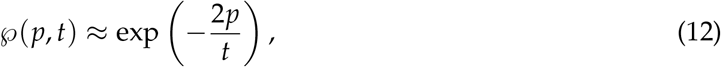

where we have replaced the boundary condition (10) with 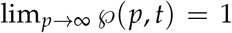. For *t* ≪ 2*p*, this probability is exponentially small, while for *t* > 2*p* it behaves like 1 — 2*p/t* (Fig. 4C).

We can use (12) to find the expected frequency of a new mutation conditional on its survival to time *t*. By the law of total probability we have

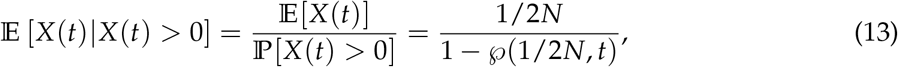

where in the last equality we used the fact that for a new neutral mutation 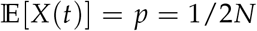. Thus, to leading order in 1/*N*, we have 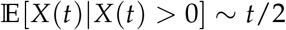.

